# Interactive webtool for analyzing drug sensitivity and resistance associated with genetic signatures of cancer cell lines

**DOI:** 10.1101/2022.08.22.504894

**Authors:** Myriam Boeschen, Diana Le Duc, Mathias Stiller, Maximilian von Laffert, Torsten Schöneberg, Susanne Horn

**Affiliations:** Rudolf Schönheimer Institute of Biochemistry, Medical Faculty, University of Leipzig, Johannisallee 30, 04103 Leipzig, Germany; Institute of Human Genetics, University of Leipzig Medical Center, Leipzig, Germany; Institute of Pathology, University of Leipzig Medical Center, 04103, Leipzig, Germany; Department of Dermatology, University Hospital Essen, University Duisburg-Essen, and German Cancer Consortium (DKTK) partner site Essen/Düsseldorf, 45122 Essen, Germany

**Keywords:** association test, candidate compound, public resource, data repository, discovery approach

## Abstract

A wide therapeutic repertoire has become available to oncologists including radio- and chemotherapy, small molecules and monoclonal antibodies. However, drug efficacy can be limited by genetic changes that allow cancer cells to escape therapy. Here, we designed a webtool that facilitates the data analysis of the Genomics of Drug Sensitivity in Cancer (GDSC) database on 265 approved compounds in association with a plethora of genetic changes documented for 1001 cell lines in the Cancer Cell Line Encyclopedia (CCLE, cBioPortal). The webtool computes odds ratios of drug resistance for a queried set of genetic alterations. It provides results on the efficacy of single or groups of compounds assigned to cellular signaling pathways. Using this webtool we replicated known genetic drivers and identified new candidate genes, germline variants, co-mutation, and pharmacogenomic modifiers of both cancer drug resistance and drug repurposing. Webtool availability: https://tools.hornlab.org/GDSC/.

**Author Summary:** Tumors develop through uncontrolled cell growth enabled by various alterations that create tumor heterogeneity. Changes of the genome and thus cancer cells cause every patient to react differently to drugs and can lead to drug resistance in cancer therapies. To overcome drug resistance, researchers focus on developing personalized therapies. Here, we provide a straightforward tool to test public *in vitro* drug sensitivity data on a range of drugs for custom analyses of genetic changes. This may inform the identification of potential drug candidates and improve our understanding of signaling pathways as we can test drug response with custom sets of genetic changes according to specific research questions. The tool and underlying code can be adapted to larger drug response datasets and other data types, e.g. metabolic data, to help structure and accommodate the increasingly large biomedical knowledge base.

## Introduction

Genetic diversity across cancers modifies the tumor’s drug responsiveness and impedes the use of widely applicable drugs without patient stratification (1). The *BRAF* V600-specific inhibition of tumor growth across various tumor entities is a prime example of personalized cancer treatment (2) and depends on the availability of diagnostic biomarkers with predictive power for drug response (3). To identify and establish those powerful decisive markers, it is crucial to associate specific genetic changes with drug response data. Complex datasets have been generated for cancer cell lines to provide information on genetic changes and drug response. The Genomics of Drug Sensitivity in Cancer Project (GDSC) and The Cancer Cell Line Encyclopedia (CCLE) represent two of such databases (4,5). While GDSC provides drug response data including bioinformatically estimated concentration-response curves, IC_50_ values as well as tested genomic associations, the CCLE dataset holds information on altered gene copy number, point mutations, mRNA levels, and gene fusions. Although public webtools were developed for ready access and analysis of the above datasets (6–10), these tools often lack the possibility to analyze a custom set of mutations or a combination of co-occurring gene alterations. Here, we designed a straightforward publicly accessible tool that combines the GDSC drug response data and genetic data collected for the CCLE samples. It enables researchers to query individual combinations of genetic alterations to assess drug sensitivity for a larger variety of cell lines. Our approach allows any combination of genetic changes in the query genes to address cellular pathways. Since different genetic changes of the same gene can mediate various functional effects, we offer custom queries for hypothesis testing and discovery of drugs for potential repurposing.

## Results and Discussion

As a proof of principle, we tested established gene-drug associations across tumor entities. Specific gene mutations are routinely used as biomarkers to guide targeted therapies, such as *BRAF* V600 mutations for the targeted MAP kinase inhibition (MAPKi) therapy of cancer patients (2). We replicated this established association with strong statistical support in our webtool. *BRAF* V600-positive cell lines were sensitive to compounds targeting the ERK-MAPK signaling pathway (OR=0.16, p=3.2e-125) such as PLX4720 (OR=0.03, p=6.2e-38) and dabrafenib (OR=0.02, p=3.87e-36, Supplementary Fig. 2, see Video 1). Of note, *BRAF* V600-mutated cell lines were resistant to compounds targeting the PI3K/mTOR pathway (OR=1.72, p=1.12e-7), EGFR pathway (OR=3.10, p=2.37e-7), and the RTK pathway (OR=1.82, p=1.47e-9, Supplementary Fig. 3A). Since RAS proteins activate the MAPK-signaling pathway (12), we expected cancer cell lines carrying one of the recurrent point mutations in *NRAS, HRAS* or *KRAS* to be sensitive to compounds targeting the ERK-MAPK signaling pathway as well. This alleged association was confirmed across entities with great statistical support (OR=0.49, p=3.71e-33, Supplementary Fig. 3B). As the role of *EGFR* and *ERBB2* overexpression and amplification in cancer is well-described (13), we expected to replicate a sensitivity to compounds inhibiting the EGFR-signaling pathway in *EGFR-* or *ERBB2-*altered cell lines. Cell lines with potentially activating changes including gene amplifications as well as high mRNA levels in either *EGFR* or *ERBB2*, or both genes were associated with sensitivity to EGFR-pathway compounds (OR=0.51 p=3.05e-8, Supplementary Fig. 4). Compounds that showed sensitivity in *EGFR/ERBB2-altered* cell lines almost exclusively target the EGFR pathway (**Fig. 2A**). Furthermore, as PTEN loss results in hyperactive PI3K signaling (14) we tested cell lines with *PTEN* deactivating changes and confirmed those to be sensitive to compounds inhibiting the PI3K/mTOR pathway (OR=0.75, p=2.94e-5, Supplementary Fig. 5A, B). Hence, we replicated several established associations of genetic changes and drug sensitivity across cancer entities, indicating the reliability of the presented webtool to detect true strong associations.

**Figure 1.**
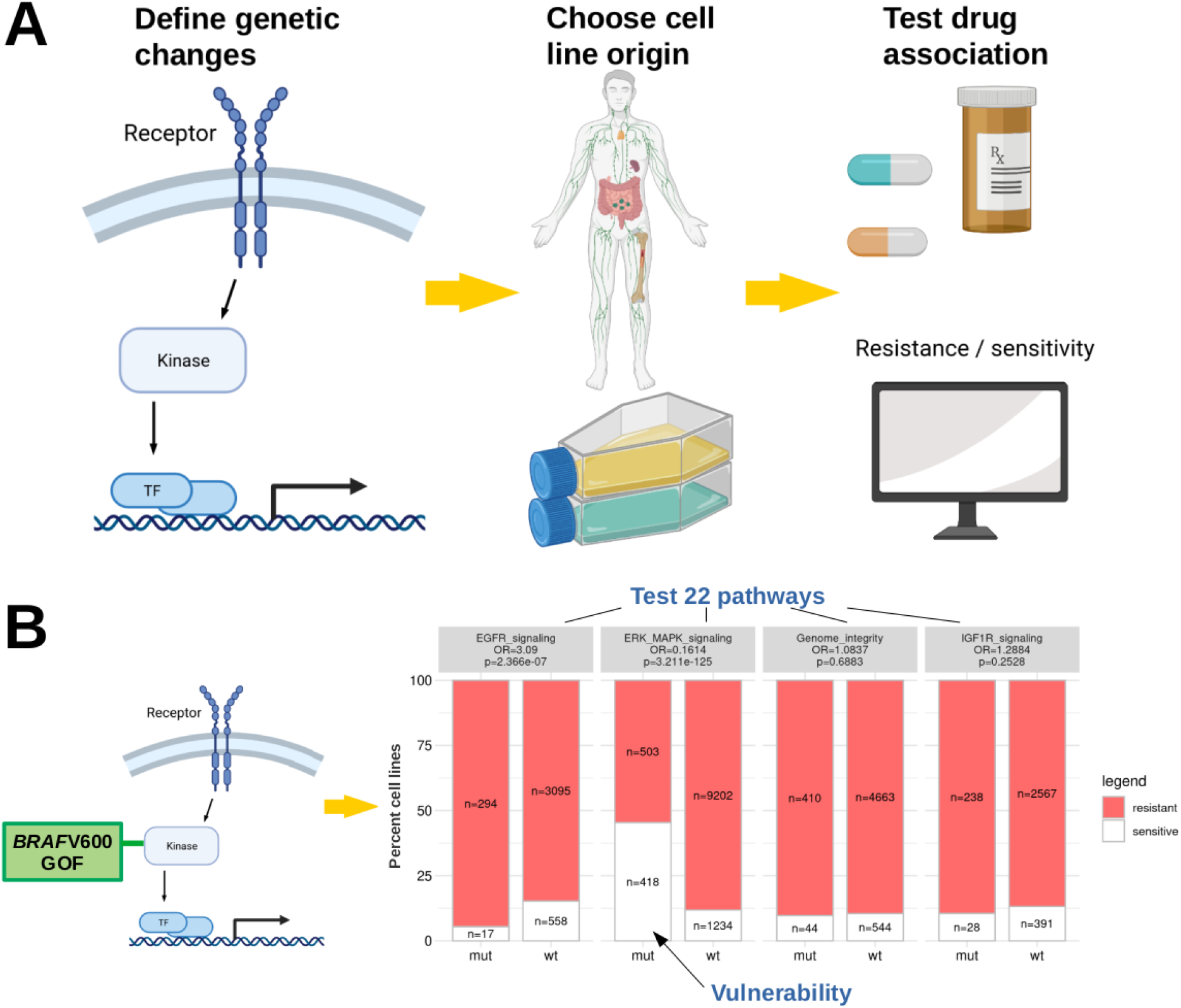
Workflow of the webtool and verification of the *BRAF*-MAPK pathway association. **(A)** Users define custom genetic changes and test across cell lines from various cancer entities. Categorization in ‘resistant’ or ‘sensitive’ from (11): cell viability measured using the CellTiter-Glo^®^ Assay, IC50 values subsequently estimated by a non-linear mixed effects model, binarization threshold for each drug. **(B)** Principle of testing the gain of function (GOF) *BRAFV600* mutated cell lines revealing moderate resistance to EGFR and strong sensitivity to ERK/MAPK pathway compounds. Odds ratios (OR) for resistance in the altered cell lines (mut) above each bar plot (Fisher’s tests). Numbers of cell lines pooled in each pathway category differ by available data indicated in the bars. Using Bonferroni correction for testing 22 pathways, p-values <0.0023 are considered significant.

**Figure 2.**
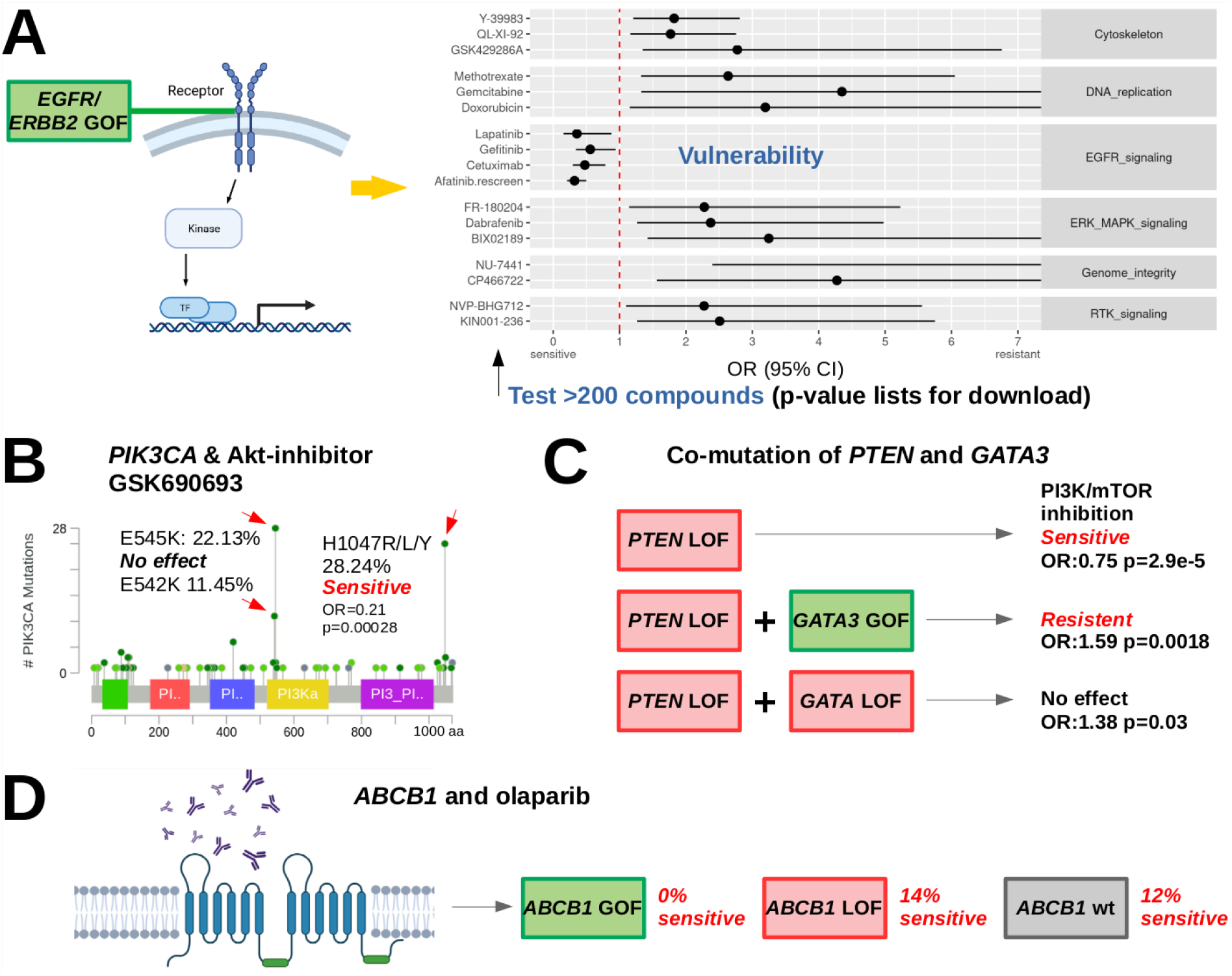
Verification of EGFR pathway association and new drug dependencies. **(A)** Compounds that show activity in *EGFR/ERBB2* activated cell lines almost exclusively belong to the EGFR pathway. Forest plot with OR>1 shows resistance of altered cell lines to drugs, OR<1 indicates sensitivity. Compounds shown if p<0.05 from Fisher’s exact tests for candidate approaches. For discovery testing 250 compounds, Bonferroni corrected p<0.0002 are considered significant. **(B)** Within frequent *PIK3CA* point mutations only H1047 mutant cell lines were sensitive to AKT-inhibitor GSK690693 (Supplementary Table 3). Frequencies and image for CCLE 2019 from www.cbioportal.org. Colored regions: protein domains. Light green: missense variants of unknown significance, dark green: driver missense mutations. **(C)** *GATA3* activation and abolished *PTEN*-related sensitivity to PI3K/mTOR inhibition. **(D)** Model of olaparib exclusion with activated *ABCB1*. GOF/LOF: gain/loss of function.

Next, we tested proposed gene-drug associations, at first the ‘BRCAness’ hypothesis (15), comparing cancer cell lines with deactivating changes in at least one of the following ‘BRCAness’ genes: *BRCA1, BRCA2, ATM, ATR, RAD51C*, and *RAD51D* with wild-types across entities. The sensitizing effect of *BRCA* deactivation for poly(ADP-ribose) polymerase inhibition (PARPi) therapy has been shown multiple times across cancer entities (16). However, we found no evidence for an increased sensitivity to the assayed PARP inhibitors (Supplementary Table 2). These results indicate that ‘BRCAness’ may not be an obligate prerequisite for benefit from PARPi in line with recent studies (17). As a second test of a previously reported association we tested PI3K/AKT/mTOR inhibitors and *PIK3CA* mutations. Somatic mutations have been reported to occur in cancer as well as pathogenic germline variants in *PIK3CA*-related overgrowth. We tested a set of 21 drugs from the group of PI3K/AKT/mTOR inhibitors and observed support for the described sensitivity of the H1047 mutant cell lines (18) for at least one inhibitor in the pathway, the AKT inhibitor GSK690693 (OR=0.21, p=0.00028 at alpha=0.002, **Fig. 2B**, Supplementary Table 3). Furthermore, *PIK3CA* E545K or E542K mutated cancer cell lines had been described to be less sensitive (18) and indeed were not associated with an apparent response to GSK690693 (OR=0.81, p=0.69). Beyond cancer, the mutational spectra of *PIK3CA*-related cancer and *PIK3CA*-related overgrowth syndromes (PROS) overlap (19). Patients with PROS showed improvements in disease symptoms following treatment with the PI3K inhibitor BYL719 and we thus tested the most frequent PROS *PIK3CA* mutations H1047, E542 and C420. Although BYL719 is not included in our drug dataset, cell lines harboring one of the 3 mutations also showed sensitivity to the Akt inhibitor GSK690693 (OR=0.28, p=0.00019). Hence, we support the notion that blocking Akt is efficient in the presence of one of the three most frequent *PIK3CA*-overgrowth mutations.

To test drug resistance of co-mutations in a large cancer entity we chose the significantly co-mutated gene pair *PTEN* and *GATA3* (co-mutation p<0.001, Supplementary Fig. 5C-E). *GATA3* expression was described to delay tumor progression and reduce Akt activation in *PTEN*-deficient mouse prostate cancer (20) and results in differential drug sensitivity in breast cancer (21). While *PTEN* deactivation was associated with the previously observed sensitivity to compounds in the PI3K/MTOR pathway, a concomitant *GATA3* activation conferred resistance (OR=1.59, p=0.0018, **Fig. 2C,** Supplementary Table 4). A concomitant *GATA3* deactivation did not yield a significant result (OR=1.38, p=0.029, significance level testing 22 pathways: 0.002).

Next, we tested putatively activating genetic changes of the pharmacogenomic modifier *ABCB1* vs. the wild-type and found resistance to compounds from various pathways (Supplementary Table 5). Olaparib efflux has been reported for advanced prostate cancer over-expressing *ABCB1* (22) and also here all of the *ABCB1* activated cell lines were resistant to olaparib (n=76, p=0.00015, **Fig. 2D**), while 12.4% of wild-type cell lines were sensitive. A deactivation of *ABCB1* in 21 cell lines had no significant association with olaparib efficacy (OR=0.82, p=0.7373).

In conclusion, we enable the swift analysis of molecular tumor heterogeneity to identify compounds for drug repurposing and prioritization in targeted therapy regimes, settings of co-mutation and multidrug resistance. The open source of our webtool is technically applicable to larger datasets with binary and continuous measures for drug response to bridge from large drug sensitivity screens to bench-side researchers.

## Materials and Methods / Design and Implementation

### Implementation

The webtool was implemented using R shiny in R version 4.0.5. To avoid duplications of 15 compounds screened twice, the screenings with higher data availability were used. Drugs were assigned to one of 22 target pathways as previously described (11). Here, all genes were analyzed across all available cell lines as the majority of cell lines were profiled for the queried alterations. Users can specify which genetic changes and cell lines shall be included in the analysis, for every gene the type of genetic changes can be selected independently (Supplementary Fig. 1). Furthermore, cell lines with missing data, which were not profiled for a specific type of alteration, can be excluded to avoid ascertainment biases in the analyses e.g. of gene fusions which have not been profiled for the majority of cell lines. Plots and data tables are provided for download.

### Statistical analysis

From the contingency tables of genetically changed and wild-type groups, odds ratios (OR) for resistance and Fisher’s exact tests (two-sided) were computed, within each pathway as well as for every single compound. Bonferroni corrected p-value thresholds are <0.0023 testing 22 pathways and p<0.0002 testing 250 drugs. For counts of zero, it was not possible to calculate OR and ratios of resistant and sensitive cell lines are shown in percent above the bar plot.

## Data availability statement

Generated code, group definitions of cell lines and resulting tables are available at https://github.com/MyriamBoeschen/Drug_Response_Tool, data sources in Supplementary Table 1.

## Authors’ Contributions

M. Boeschen: Conceptualization, data curation, software, formal analysis, validation, investigation, visualization, methodology, writing-original draft. D. Le Duc: Conceptualization, writing-review and editing. M. Stiller: Conceptualization, writing-review and editing. M. von Laffert: Conceptualization, writing-review and editingConceptualization, writing-review and editing. T. Schöneberg: Conceptualization, supervision, funding acquisition, writing-review and editing. S. Horn: Conceptualization, methodology, formal analysis, validation, investigation, visualization, supervision, project administration, funding acquisition, writing-review and editing

## Acknowledgments

We thank Udo Stenzel for advice on the rationale and implementation of this webtool. This work was funded by the Deutsche Forschungsgemeinschaft (DFG, German Research Foundation) HO 6389/2-2, ‘KFO 337’ - 405344257 to SH and SFB 1052 Project B10 to DLD. The funders were not involved in study design, data collection and analysis, decision to publish, or preparation of the manuscript.

## Supplementary data

Supplementary File 1. PDF containing Supplementary Figures 1-5 and legends in the order cited in the manuscript.

Supplementary File 2. Open office document containing Supplementary Tables 1-5 and legends in the order cited in the manuscript.

